# Crystalline silica exposure causes latent biomechanical abnormalities along the cardiopulmonary axis

**DOI:** 10.64898/2026.10.20.760460

**Authors:** Meredith S. Gwin, Jack Di Palo, Anthony A. Fung, Rira Choi, Pete H. Gueldner, Alexandria Regan, Aishani Nagamalla, Asad A. Syed, Theodore Sandow, Ying Sun, Sydney L. Bluestein, Buqu Hu, Yosep Shin, Keshav Harpavat, Clemente Britto, P. Kent Langston, Jose Gomez, Xinran Liu, Nicole Guerrera, Jonathan L. Koff, Xiting Yan, Jason M. Szafron, Susan M. Majka, Rong Fan, Zachary M. Harris, Edward P. Manning

## Abstract

**Rationale:** Crystalline silica exposure is a major global health burden linked to chronic lung disease and cardiopulmonary morbidity, but early lung-vascular-cardiac abnormalities preceding overt functional decline are poorly defined.

**Objectives:** To determine whether a single crystalline silica exposure produces subclinical abnormalities across the integrated axis of cardiopulmonary response (lung→pulmonary artery→right ventricle) and identify associated sex-dependent physiologic patterns and molecular programs.

**Methods:** Male and female mice received a single endotracheal instillation of crystalline silica or vehicle (saline). At 28 days, assessments included voluntary wheel running, diffusion capacity, pulmonary function testing, bronchoalveolar lavage analysis, lung histology and electron microscopy, *ex vivo* proximal pulmonary artery biomechanics and microstructural second harmonic generation imaging, right ventricular pressure, transthoracic echocardiography, and bulk RNA sequencing of lung and right ventricle.

**Results:** Despite recovery of body weight and preserved voluntary exercise, silica-exposed mice demonstrated reduced diffusion capacity, ultrastructural thickening, and disruption of the alveolar-capillary barrier. Bronchoalveolar lavage revealed persistent inflammation with elevated macrophages, lymphocytes, neutrophils, and protein, alongside retained particulate and surfactant-rich debris (silicoproteinosis). Lung mechanics were sex-dependent; females showed a parenchymal-dominant phenotype and males showed increased airway resistance. Elevated pulmonary circulatory pressures caused increased functional stiffness and reduced distensibility of the proximal pulmonary arteries. Right ventricular free wall thickening was noted in females. Lung transcriptomics indicated persistent immune activation and extracellular matrix remodeling, while the right ventricle demonstrated metabolic reprogramming with downregulation of structural and reparative programs.

**Conclusions:** A single crystalline silica exposure induces a subclinical, multi-compartment cardiopulmonary phenotype characterized by persistent alveolar-capillary injury, pulmonary vascular stiffening, increased right ventricular afterload, sex-dependent pulmonary mechanics, and divergent organ-level transcriptomic responses.

## INTRODUCTION

Increasing numbers of veterans and other exposed populations are being diagnosed with deployment-related respiratory disorders linked to burn pits, particulate matter, dust, and other airborne toxicants, highlighting an urgent need to elucidate system-level mechanisms preceding overt decline following toxic inhalant exposure (*1, 2*). Silica exposure is primarily recognized as a chronic occupational exposure in the mining, quarrying, and construction sectors as well as wildland firefighters exposed to airborne particulate matter in wildfire emissions (*4–8*). In addition, over two million post-9/11 veterans deployed to Southwest Asia (SWA) were exposed to the silica dust (intermittent high burdens during sandstorms and continuous low levels caused by agitated sand secondary to vehicle traffic and explosions) leading to a significant population of exposed individuals (6–8). Lung biopsies from veterans deployed to SWA revealed significantly higher silica particle counts compared to those from non-servicemembers (9). Despite up to 16% of deployed veterans reporting chronic adverse cardiopulmonary symptoms (dyspnea, cough, decreased exercise tolerance), clinical diagnoses to explain them are less forthcoming (10–13). Deployment to SWA is increasingly recognized as a predictor for poor pulmonary health outcomes (*9–11*). Millions of workers, wildland firefighters, and post-9/11 veterans face compounding or cumulative crystalline silica inhalation exposures that drive progressive, irreversible chronic lung injury. The lack of advanced phenotyping to define exposure-induced mechanisms of lung injury and to link those mechanisms to the cardiopulmonary symptoms experienced by individuals with deployment- and occupational-related disease creates a critical gap in diagnosis and treatment.

Silica-rich dust has been linked as causative to respiratory symptoms and chronic diagnoses with documented airway and parenchymal abnormalities (e.g., small-airway disease/constrictive bronchiolitis, inflammatory remodeling, and imaging/biopsy changes) (*3*). To date, a central mechanism underlying the clinical features of silica exposure and pathogenesis of latent cardiopulmonary symptoms (dyspnea and exertional intolerance) is lacking due in part to the frequent latency between exposure and overt disease, complicated by non-specific diagnostic testing, which delays recognition and treatment.

This gap persists for two interconnected reasons. First, exertional dyspnea is a property of the integrated cardiopulmonary system, one that requires coordinated, dynamic measurements across multiple physiologic domains. Clinically meaningful impairment is frequently unmasked only under physiologic stress and remains invisible to standard resting tests (*12*). Second, deployment- and occupational-relevant particle and toxicant injury can plausibly initiate chronic, self-propagating remodeling cascades, encompassing immune dysregulation, fibroproliferation, microvascular pruning, and arterial stiffening that evolve subclinically until cardiopulmonary reserve is sufficiently eroded to produce overt symptoms (*13–18*).

Closing this gap demands a systems-level investigation that couples lung function to the right ventricle (RV) via the pulmonary artery (PA) (the lung-PA-RV axis) that simultaneously characterizes, pulmonary vascular remodeling and mechanics, and RV function. Prioritizing exertional physiologic assessment (CPET, 6MWT) and exertional oximetry over resting spirometry alone may enable the detection of early vascular and RV alterations within the "normal test" window (*19*). This links subclinical structural changes to patient-reported symptoms, predicts subsequent functional decline, and supports earlier risk stratification and targeted intervention in alignment with the real-world Veterans Health Administration (VHA) phenotype (*20*).

Critically, studying inhalation exposures across the lung→PA→RV continuum provides the mechanistic resolution needed to identify drivers of adverse respiratory symptoms that elude conventional clinical diagnosis. For these reasons, we investigated silica exposure across the integrated cardiopulmonary system in a murine model, incorporating functional endpoints, including pulmonary function testing (PFT), transthoracic echocardiography, and voluntary exercise; and mechanistically informative measures that include and extend beyond standard clinical markers, such as diffusing factor for carbon monoxide (DFCO) and pulmonary artery mechanical testing. We hypothesized that a single, high-burden endotracheal installation of crystalline silica produces an inflammatory response in the lung that drives PA stiffening and RV compensatory changes. This work will aid in defining how the functional and ultrastructural mechanisms of silica exposure impact the comprehensive cardiopulmonary system while also establishing a repeatable model for future respiratory exposure studies.

## METHODS

### Mice

Animal experiments were designed, conducted, and reported in accordance with the ARRIVE 2.0 guidelines to promote rigor, transparency, and reproducibility in preclinical research(*21*). It was approved by the Yale University Institutional Animal Care and Use Committee. Female and male C57BL/6J WT mice aged 13-15 weeks old were used for these studies.

### Silica

Silica was purchased from naturalpigments.com (SKU# 510-155LC). Median particle size is 1.6 μm with 96.3% of particles being finer than 5 μm.

### Exposure

Crystalline silica (0.4 g/kg of bodyweight) was endotracheally administered suspended in a 0.9% saline solution vehicle. Control mice were administered only the vehicle. The total volume of solution administered was 3.33 μL/g of bodyweight.

### Voluntary Exercise

Daily running distance of mice was measured using in-cage running wheels (Actimetrics Model ACE-557-WLP) with ClockLab Data Collection Software from Lafayette Instrument. The running wheels are designed to not interfere with normal housing conditions. The mice were placed in the cages with running wheels for a total of three days, and their voluntary exercise was reported as the daily run distance (24-hour period) averaged over the three days. The mice were singly housed during these experiments and had access to food and water *ad libitum*.

### Transthoracic Echocardiography

Noninvasive investigation of cardiac function was performed in mice using transthoracic echocardiograms under light anesthesia (1.5% isoflurane) while maintaining physiologic temperature (*22*).

### Lung Measurements

Adequate depth of anesthesia was confirmed by absence of a toe-pinch withdrawal reflex, and supplemental urethane was administered as needed. DFCO was then measured using a gas chromatograph (Inficon Micro GC Fusion) (*23*). After DFCO assessment, pulmonary mechanics were measured using the FlexiVent system. Upon completion of mechanical testing, bronchoalveolar lavage (BAL) fluid was collected.

### PA Biomechanical Measurement and Analysis

We define proximal PA as the extralobar PA and distal PAs as those in the lung. We performed *ex vivo* biaxial mechanical testing of proximal PAs using a custom-made, computer-controlled device as previously described (*22, 24*). Results were fit using the Four-fiber family constitutive model for arterial walls as previously described, quantifying stiffness, distensibility and other material and biomechanical properties across physiologic loading ranges (*25, 26*).

### RNA sequencing

RNA was extracted from the flash-frozen samples of lung and RV, stored at –80 °C using miRNeasy Micro kits (Qiagen, 217084) following manufacturer’s protocol. Purified total RNA was submitted to Plasmidsaurus (Eugene, OR) for 3′-end counting RNA sequencing.

### Statistics

Data were analyzed by Welch’s t-test unless otherwise stated. All analyses were made using GraphPad Prism (11.1.0). Data is presented as ± standard error from the mean (± SEM). Significance was defined as a p-value < .05, except for RNA sequencing data, where a Benjamini-Hochberg adjusted p-value (FDR) < .05 was used to account for multiple hypothesis testing.

### Histology

Tissues were fixed in 10% neutrally buffered formalin (NBF) overnight, and then moved to a 70% ethanol solution and kept at 4 °C. The left lung specifically was inflated to 20 cmH_2_O of pressure prior to and during fixation to allow for morphological analysis.

### Histological and physiologic biomarker classification and analysis

See supplemental.

Detailed methods for each experiment can be found in Supplementary Material.

## RESULTS

### Silica exposure causes subclinical cardiopulmonary dysfunction

Occupational silica exposure remains a significant global health concern, yet the subclinical cardiopulmonary consequences that precede overt clinical disease are poorly understood. While silicosis is classically defined by progressive pulmonary fibrosis and impaired lung function, early pathophysiological changes at the alveolar-capillary interface that occur well before functional decline are present (Figure 1A) and may be clinically apparent if sought. Therefore, understanding these subclinical changes is critical, as they may represent a window for early intervention and could help explain the disproportionate cardiovascular morbidity observed in silica-exposed workers (*27–29*).

**Figure 1.**
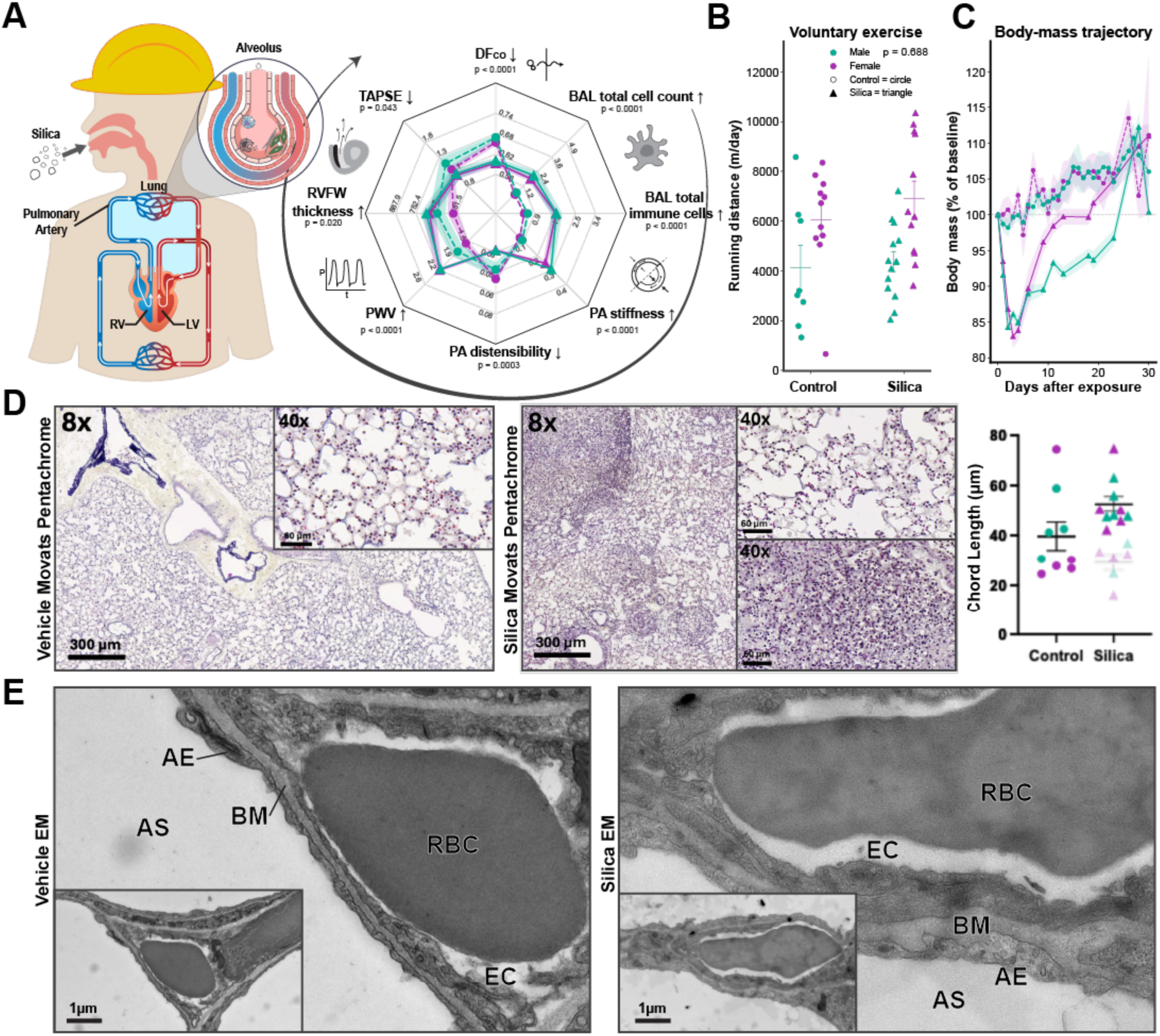
Exposure to silica causes insidious cardiopulmonary dysfunction. Hypothesized propagation of silica exposure pathophysiology manifests at multiple points in the pulmonary circuit, with multiple physiologic biomarkers expressed as means with 1-standard deviation error bands (panel A). Cardiopulmonary function and body weight are preserved 28 days after silica exposure (panels B and C, respectively) despite impaired gas exchange in the lung (decreased diffusion capacity, panel C *P* < .0001). Alveolar size increased in regions of the lung without infiltration (opaque data points) and decreased in regions of the lung where infiltration was present (translucent data points), though these trends were not significant (panel D). Representative image of control female mouse (exposed to vehicle only, 0.9% saline, panel D left image) is exhibited in comparison with a representative image of a lung from a female mouse exposed to 0.4g/kg silica in 0.9% saline 28 days earlier (panel D right image). The exposed mouse lung exhibits heterogeneous regions comprising areas of alveolar enlargement (top inset) and monocytic-predominant infiltration (bottom inset). We found significant thickening and disruption of the shared basement membrane (BM) between capillary endothelial cells (EC) and epithelial cells (AE) in mice exposed to silica (panel E left image, representative electron micrograph of control mouse; panel E right image, exposed mouse; AS = alveolar space). Data is represented as mean (± SEM). n = 6 – 26 per group.

28 days after exposure to crystalline silica (0.4 g/kg of bodyweight) via endotracheal instillation, mice maintained baseline cardiopulmonary exercise tolerance, as measured by voluntary running distance (Figure 1B). Evidence of acute injury was observed as mice lost ∼16% of their body mass within the first 4 days of exposure followed by recovery to pre-exposure body weight by day 28 (Figure 1C). 28 days after exposure, despite apparent recovery we found that DFCO was significantly reduced in silica-exposed mice compared to controls (Figure 1A, *P* < .0001), quantitating persistent impairment of lung gas exchange across the alveolar-capillary membrane. The size of alveoli tended to be larger in non-infiltrated regions of the lung and smaller in infiltrated regions quantitated by mean chord length (Figure 1D). Ultrastructural transmission electron microscopy analysis of the alveolar-capillary unit highlighted evidence of thickened endothelial-epithelial shared basement membrane (Figure 1E), further supporting a mechanism of impaired diffusion quantitated by DFCO (Figure 1A).

### Alveolar-capillary disruption characterized by inflammation and leak are present 28 days after silica exposure

The development of silicosis and other silica-related lung diseases is fundamentally driven by a persistent inflammatory response and the subsequent breakdown of the delicate alveolar-capillary membrane. To understand how a single silica exposure leads to sustained lung dysfunction, markers of lung inflammation and breakdown of the alveolar-capillary membrane were examined in silica-exposed mice compared with saline controls. BAL was collected 28-days post-silica exposure and analyzed for cell count, cell differential, and total protein. Silica-exposed mice showed a significant increase in BAL total cell count compared with saline controls (Figure 2A, *P* < .0001). BAL total macrophages, lymphocytes, and neutrophils were increased in silica-exposed mice compared with saline controls (Figure 2B-D; *P* = .0003, *P* < .0001, and *P* < .0001, respectively). BAL cell differential demonstrated significantly decreased percentage of macrophages and significantly increased percentages of lymphocytes and neutrophils compared to saline controls (Supplemental Figure S1). There was also a significant increase in the BAL total protein in silica-exposed mice compared to saline controls (Figure 2E, *P* < .0001) indicative of alveolar-capillary barrier leak. We also found foamy macrophages in the BAL as well as isolated silica particles on histology in alveoli of mice exposed to silica (Figure 2F). We verified the presence of macrophages and proteinaceous debris in the airway using electron microscopy, including non-degraded surfactant and other debris (Figure 2G, Supplemental Figure S2). Together, these data demonstrate sustained lung damage 28 days after a single silica exposure that associates with endothelial-epithelial cell barrier dysfunction.

**Figure 2.**
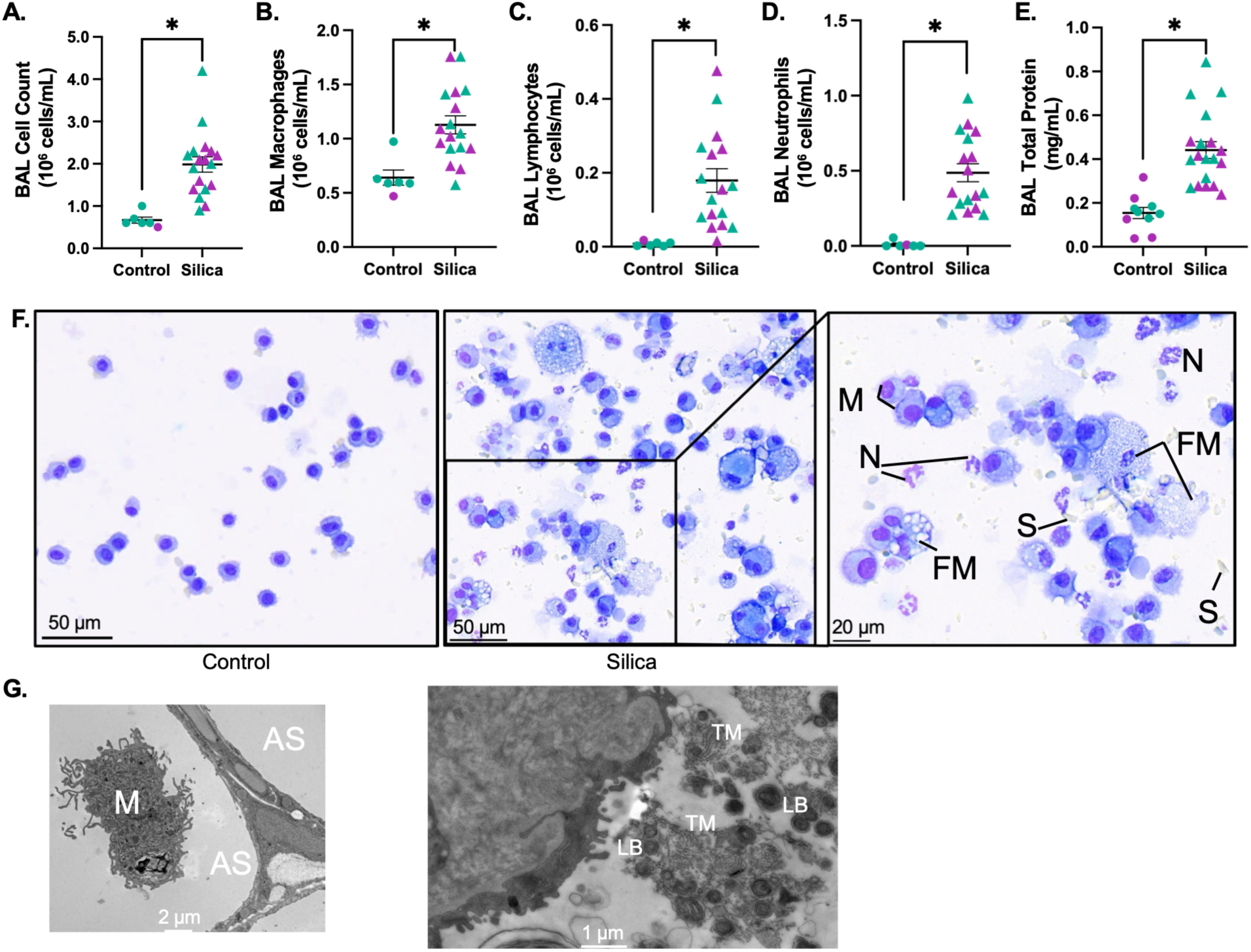
Pulmonary inflammation and alveolar-capillary disruption are present 28 days after silica exposure. BAL collected 28-days post-exposure to silica showed a significant increase in total cell count compared with saline controls (panel A, *P* < .0001). BAL total macrophages, lymphocytes, and neutrophils were increased in silica-exposed mice compared with saline controls (panels B-D) (B, *P* = .0003; C, *P* < .0001; D, *P* < .0001). There was a significant increase in the BAL total protein in silica-exposed mice compared with saline controls (panel E, *P* < .0001). BAL from controls revealed alveolar macrophages (panel F, left image). Accumulated foamy macrophages (FM), silica particles (S), neutrophils (N), and macrophages (M) were present in BAL of mice exposed to silica 28 days earlier (panel F, middle and right images). We confirmed the presence of macrophages (M) in alveolar space (AS) of control mice on electron microscopy (panel G, left image) and, lamellar bodies (LB), and tubular myelin (TM) in the alveolar space of mice exposed to silica (panel G, right image). Data is represented as mean (± SEM).

### Silica exposure produces sex-dependent pulmonary phenotypes, with parenchymal changes predominating in females and airway disease predominating in males

To assess the effects of silica-exposure on biomechanical functions of the lung, we performed pulmonary function testing. We found sex-dependent differences in lung function. Female mice exposed to silica demonstrated biomechanical changes implicating lung parenchyma, including increased area between the loading and unloading pressure-volume curves, termed hysteresis (Figure 3A, *P* = .0133); decreased tissue elastance (Figure 3B, *P* = .0100); increased parenchymal heterogeneity as measured hysteresivity (Figure 3C, *P* = .0408); and increased inspiratory capacity (Figure 3D, *P* = .0010). Male mice exposed to silica demonstrated biomechanical changes implicating conducting airways, characterized by increased Newtonian/airway resistance (Figure 3E, *P* = .0048). Between silica-exposed and control groups, parameters typically associated with parenchymal changes were not statistically different in the male cohort, while airway resistance was not statistically different in the female cohort (Figure 3A-E). While mice exposed to silica had increased debris in the parenchyma, we found that male mice had a higher proportion filled with debris than female mice (male = 21.6%, female = 16.1%, Supplemental Figure S3). On average, adult male mice have greater body mass than female mice, and it would be reasonable to suppose a particle of fixed diameter may cause biomechanical changes that are size-dependent. Therefore, we performed allometric analysis and found that these responses were independent of body mass and size (*30*). Therefore, our results are likely a reflection of sex as a biological variable rather than body mass (Supplemental Figure S4). Silica exposure causes a multinodular, monocytic predominant, peribronchiolar infiltrative process (Figure 3F). Ultrastructural analysis demonstrated evidence of proteinaceous debris, and non-degraded surfactant within alveoli and airways of exposed mice (Supplemental Figure S2). This closely resembles silicoproteinosis as described in humans during acute lung injury to silica exposure (*31, 32*).

**Figure 3.**
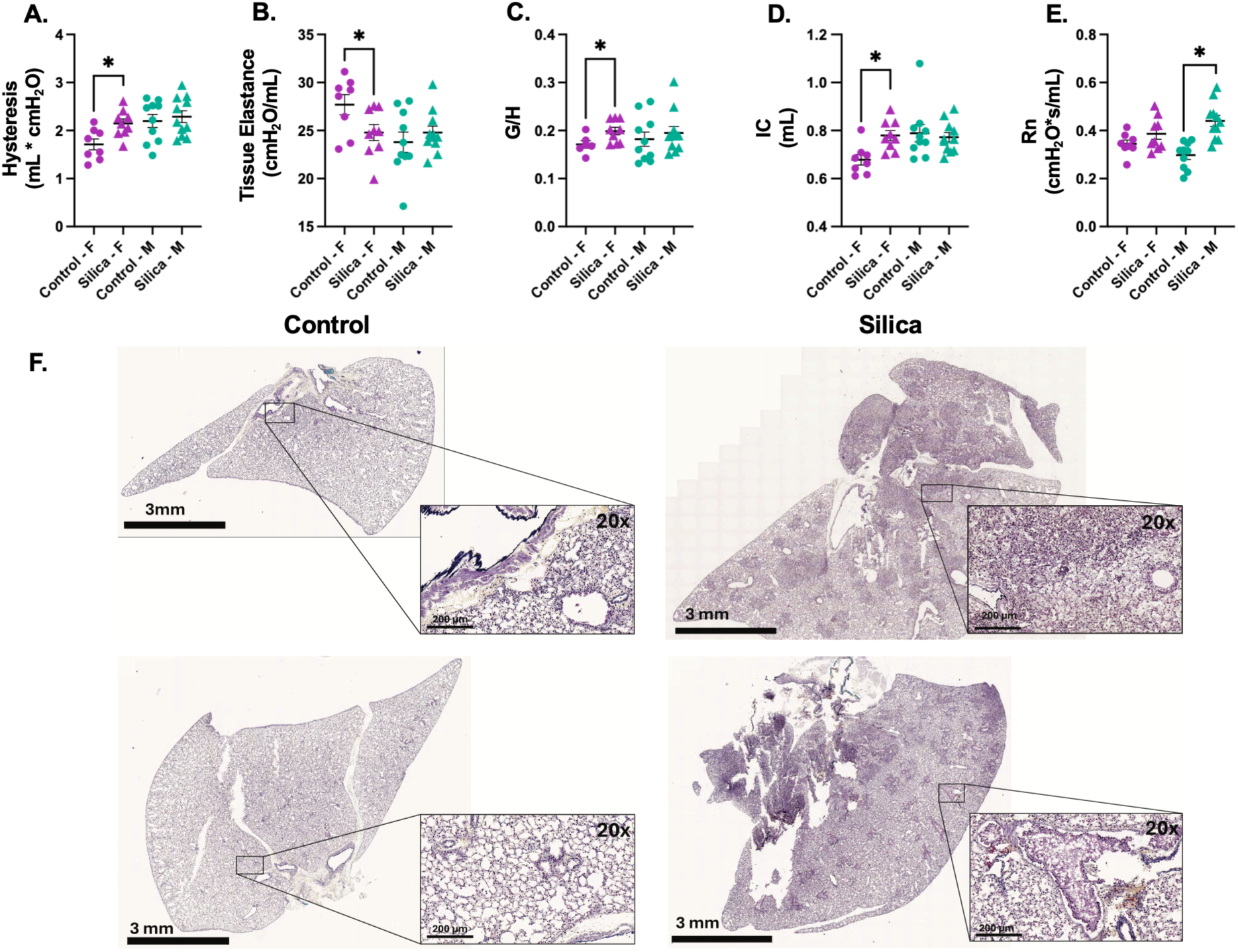
Silica exposure produces sex-dependent pulmonary phenotypes, with parenchymal disease predominating in females and airway disease predominating in males. To assess the effects of persistent silica and inflammatory cells on biomechanical functions of the lung, we performed pulmonary function testing. We found sex-dependent differences in lung function. Female mice exposed to silica demonstrated biomechanical changes implicating lung parenchyma, including increased hysteresis (panel A; *P* = .0133), decreased tissue elastance (panel B; *P* = .0100), increased hysteresivity (panel C; *P* = .0408), and increased inspiratory capacity (panel D; *P* = .0010). Male mice exhibited a predominantly airway phenotype, characterized by increased airway resistance (panel E; *P* = .0048). Representative histology reveals a picture of multinodular, monocytic predominant, peribronchiolar infiltrative process (panel F). Data is represented as mean (± SEM). n = 8-11 per group.

Pulmonary function testing demonstrated a clear sex-dependent physiologic response to silica exposure. Female mice developed a pattern most consistent with parenchymal dysfunction, reflecting altered tissue mechanics, greater energy dissipation during breathing, and loss of normal distal lung/alveolar integrity. In contrast, male mice exhibited a phenotype dominated by conducting airway dysfunction, with increased airway resistance consistent with airway narrowing or obstruction rather than primary parenchymal stiffening. These divergent responses were independent of body mass, implicating sex as a key biological determinant of the pulmonary response to silica. Histologic and ultrastructural analyses supported these functional findings, demonstrating peribronchiolar inflammatory infiltrates and retained silica with proteinaceous debris and non-degraded surfactant within both alveolar and airway compartments.

### Silica-induced lung injury is associated with a pressure-mediated mechanism of pulmonary vascular maladaptive remodeling 28 days after exposure

We analyzed geometry of the distal (intralobar) PAs in mice exposed to vehicle or silica. In mice exposed to silica, the unloaded wall thickness and the inner diameter of the vessels were unchanged compared to controls. However, there was a trend toward intimal thickening of the distal PAs in female mice suggesting remodeling of the distal, intralobar PA vessels (Supplemental Figure S5, *P* = .0631). Thickening of the micro and distal vasculature is associated with increased pulmonary vascular resistance. *In vivo* RV pressure tracings revealed the increased pulmonary circulatory pressures in silica-exposed mice (Figure 4A) and confirmed previously reported findings (*33*).

**Figure 4.**
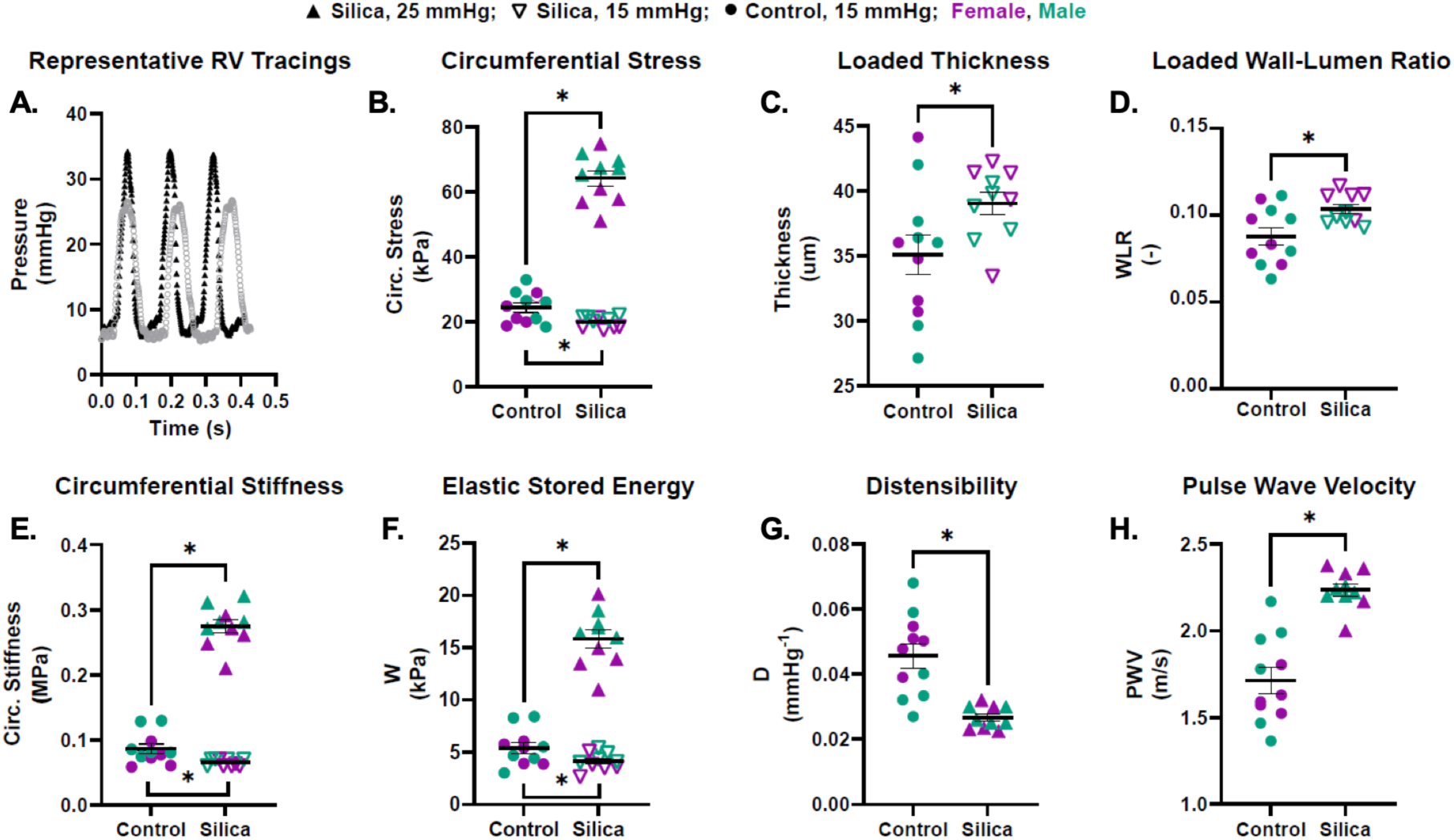
Silica-induced lung injury is associated with pulmonary vascular maladaptive remodeling 28 days after exposure. In our prior work, we found that lung injury due to hypoxic exposure was associated with increased stiffness of the proximal pulmonary arteries of mice. Therefore, we performed *ex vivo* biomechanical testing of the proximal pulmonary artery at 28 days following exposure to crystalline silica. We found increased pulmonary circulatory pressures in mice exposed to silica (representative tracings in panel A). We found that proximal pulmonary arteries in mice exposed to silica compared to controls exhibited significantly increased circumferential stress at in vivo pressures (panel B; *P* < .0001). At a common pressure of 15 mmHg, the silica-exposed mice experienced lower circumferential stress than controls (panel B; *P* = .0156), consistent with geometric remodeling to offload stress. At 15 mmHg, the proximal PAs of silica-exposed mice were thicker (panel C; *P* = .0426), driving an increase in the wall-to-lumen ratio (panel D; *P* = .0153). At in vivo pressures, proximal PAs of silica-exposed mice had greater circumferential stiffness (panel E; *P* < .0001), and a pressure-driven increase in elastic stored energy (panel F; *P* < .0001). Decreased distensibility (panel G; *P* = .0005) associates with greater pulsatile afterload burden placed on the RV, while increased pulse wave velocity (panel H; *P* < .0001) associates with greater penetration of the pulmonary vascular bed. Data is represented as mean (± SEM). n = 10-11 per group.

Given our focus on the lung-PA-RV axis, we also analyzed proximal PA mechanics due to its crucial role in modulating the pulsatile component of RV afterload and its association with RV and lung function decline (*22, 24, 34–36*).

We performed *ex vivo* biomechanical testing of the proximal PA at 28 days post exposure to crystalline silica (*33*). Comparison of PA biomechanical properties was performed at *in vivo* mean PA pressures (15 mmHg for control mice, and 25 mmHg for silica mice) as well as at a common mean PA pressure (15 mmHg for both groups). These comparisons enable us to differentiate between changes in mechanical properties resulting from differences in pressure versus those resulting from arterial wall remodeling. Solid data points in Figure 4 are *in vivo* pressure mechanical values; hollow data points are at common pressure.

The increased pulmonary circulatory pressures distend the PA and increase circumferential stress in the arterial wall (Figure 4B). Specifically, the arterial wall significantly thickens after exposure to silica (Figure 4C), significantly increasing its wall-to-lumen ratio (Figure 4D) which offloads stress from the arterial wall. Using two-photon microscopy, we found a trend toward adventitial thickening of silica-exposed mice (Supplemental Figure S6).

Despite the offloading of wall stress by geometric remodeling, increased *in vivo* pressure leads to significantly increased circumferential stiffness in the silica group (Figure 4E). Elastic stored energy at *in vivo* pressures was greater in the silica group (Figure 4F), an effect of *in vivo* pressure for the silica-exposed mice being higher than the control. Proximal PA distensibility was reduced in the silica group (Figure 4G), indicating diminished capacity to buffer pulsatile flow. Correspondingly, pulse wave velocity was increased (Figure 4H), consistent with faster pulsatile transmission along the proximal PA. Together these reflect a conduit that accommodates less volumetric change per beat, reflects increased pressure pulse waves more rapidly toward the RV, and penetrates forward pulse waves deeper into the distal bed of the pulmonary vasculature. This contributes to increased RV afterload and potential impaired gas exchange due to increased endothelial cell shear stress, respectively. Additional biomechanical properties are listed in Supplemental Tables S1 – S3.

### Silica exposure associates with signs of right ventricular compensation to elevated pressures in the pulmonary circulation

The increase in RV pressure (Figure 4A) associates with increased RV free wall thickness in female mice (Figure 5A, *P* = .02). The RV cavity trended toward increased size (Figure 5B, *P* = .13) and contractility trended down as measured by TAPSE (Figure 5C, *P* = .1010). Representative histology images of control and silica treated mice hearts are provided (Figure 5D).

**Figure 5.**
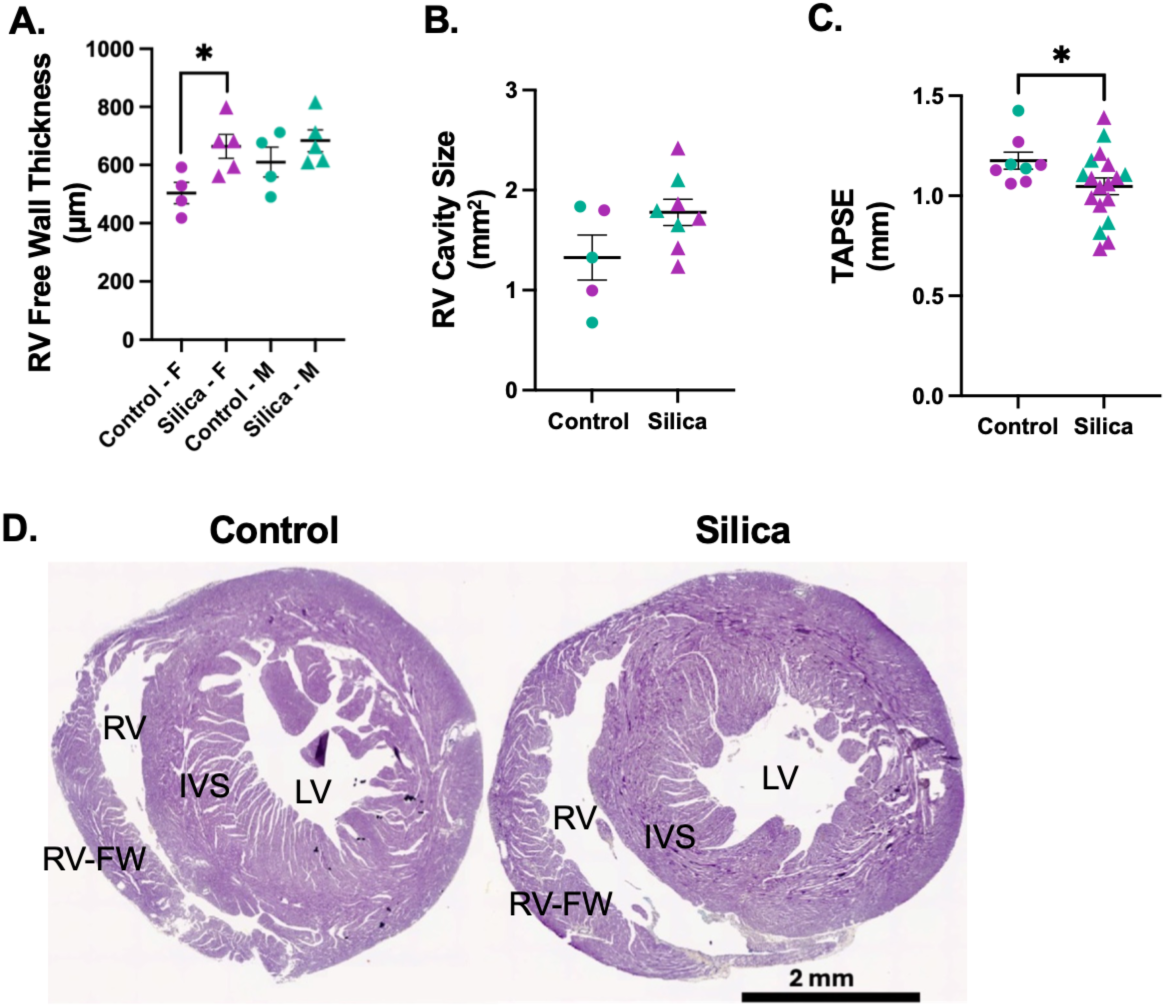
Silica exposure associates with signs of right ventricular compensation to elevated pressures in the pulmonary circulation. We observed an increase in RV pressure in mice exposed to silica 28 days earlier compared to controls (Figure 4A). The increase in RV pressure associates with significantly increased RV free wall thickness in female mice (Panel A; Female: silica 663.9μm versus control 503.8μm, *P* = .0233; Male: silica 683.4μm versus control 609.8μm, *P* = .2929). The RV cavity trended toward increased area in exposed mice (Panel B; Female: silica 1.73 mm^2^ versus control 1.40 mm^2^, *P* = .5515; Male: silica 1.85 mm^2^ versus control 1.28 mm^2^, *P* = .2250). A decrease in tricuspid annular plane systolic excursion (TAPSE) was observed in silica-exposed mice (Panel C; *P* = .0434), consistent with impaired RV function. Representative images of hearts from female hearts (Panel D). Data is represented as mean (± SEM). n = 4-12 per group. RV = right ventricle, LV = left ventricle, IVS = interventricular septum, RV-FW = right ventricle free wall.

### Transcriptomic analysis of the lung and RV tissue at 28 days post-exposure reveals divergent adaptive organ-level transcriptomic responses to silica-induced lung injury

Transcriptomic profiling of lung and RV at 28 days was done to capture transcriptomic and molecular pathways after the immediate response to silica injury, connect observed physiologic abnormalities to underlying mechanisms by which lung injury drives secondary adaptive remodeling response in the RV via the lung–heart axis (e.g., vascular injury, hypoxemia, increased RV afterload). Comparing organs distinguishes local pulmonary injury pathways from downstream cardiac adaptation and helps identify candidate biomarkers and therapeutic targets relevant to early human disease progression.

Bulk RNA sequencing of lung and RV tissue of silica-exposed mice was compared to saline controls. Silica-induced lung injury elicited an inflammatory transcriptomic response in the lung, while the right ventricle underwent a compensatory metabolic shift toward lipid metabolism.

In lungs of silica-exposed mice (Figure 6A), upregulated differentially expressed genes (DEGs) include *Tlr2, Clec7a, Clec4n*, and *Ctsd*, which were associated with damage-associated molecular programs (DAMP) (*37, 38*). Saa3 associates with a TLR2-driver positive feedback loop that may mechanistically explain the persistence of NF-kB and STAT3 signaling. *Gpnmb* and *Chi3l1* associate with dominant macrophage-driven pro-fibrotic state rather than a resolving reparation phenotype. *Mmp12* and *Ltbp2* also associate macrophage activation and ECM remodeling. The co-enrichment of interferon alpha, interferon gamma, IL-6/JAK/STAT3, and inflammatory response pathways alongside downregulation of Kras and myogenesis programs at 28 days post-silica exposure defines a transcriptomic state of persistent, maladaptive immune-driven inflammation (Figure 6B). This pattern is consistent with a maladaptive inflammatory-fibrotic transition. Ongoing immune-mediated tissue injury is not offset by compensatory epithelial or structural regeneration. The prominence of IL-6/STAT3 and interferon signaling, which we previously found to activate perivascular macrophages and adventitial fibroblast, provides a plausible mechanistic link between the lung parenchymal inflammatory response and maladaptive pulmonary vascular stiffening observed in this model. This suggests that perivascular macrophage-driven vascular maladaptive remodeling may be a downstream consequence of non-resolving silica-induced lung inflammation.

**Figure 6.**
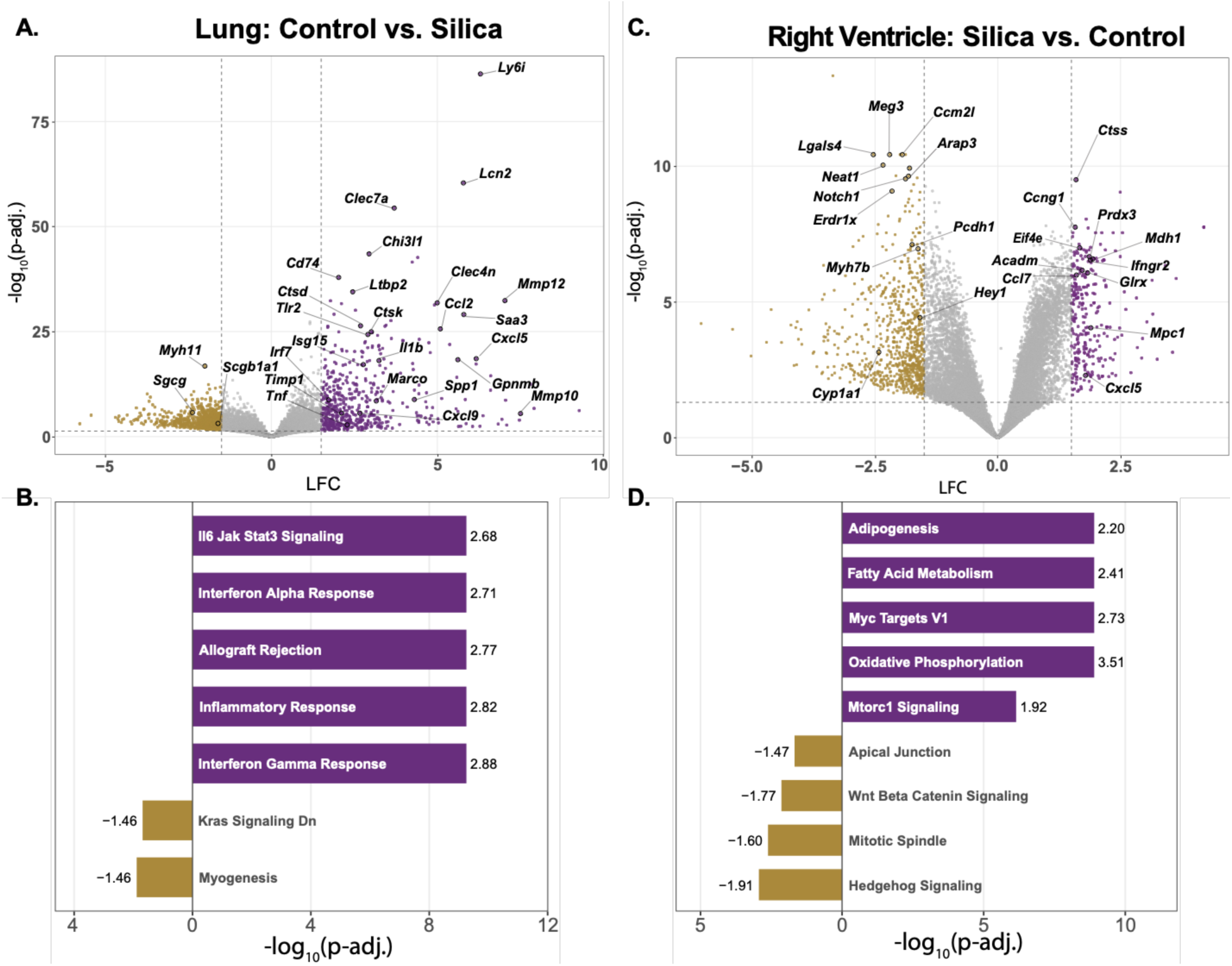
Bulk RNA sequencing of lung and right ventricular (RV) tissue at 28 days post-exposure reveals divergent organ-level transcriptomic responses to silica-induced lung injury. In the lung of silica-exposed mice (Panel A-B), upregulated differentially expressed genes (DEGs) include TLR2, Clec7a, Clec4n, and Ctsd, which are associated with damage-associated molecular programs (DAMP). Saa3 associates with a TLR2-driver positive feedback loop that may mechanistically explain the persistence of NF-kB and STAT3 signaling. Gpnmb and Chi3l1 associate with dominant macrophage-driven pro-fibrotic state rather than a resolving reparation phenotype. Mmp12 and Ltbp2 also associate macrophage activation and ECM remodeling. Gene set enrichment analysis (GSEA) showed upregulation of the interferon alpha, interferon gamma, and IL-6/JAK/STAT3 proinflammatory pathways and downregulation of Kras and myogenesis programs. In the RV of silica-exposed mice (Panel C-D), decreased DEGs, including Myh7b, Meg3, Neat1, and Ccm2l, associate with sarcomeric/cytoskeletal remodeling under hemodynamic stress while simultaneously explaining mTORC1 upregulation, Wnt loss, and fibrosis. Decreased Lgals4 and Erdr1 associate with cardiomyocyte junctional alterations, increased metabolic stress, and impaired nutrient sensing. The RV’s distinctive metabolic shift is seen in upregulation of lipid metabolism pathways.

In RV of silica-exposed mice (Figure 6C), decreased DEGs, including *Myh7b, Meg3, Neat1*, and *Ccm2l*, associate with sarcomeric/cytoskeletal remodeling under hemodynamic stress simultaneously explains mTORC1 upregulation, Wnt loss, and fibrosis. Decreased *Lgals4* and *Erdr1* associate with cardiomyocyte junctional alterations, increased metabolic stress, and impaired nutrient sensing. These combinations suggest the cytoprotective capacity of RV cardiomyocytes is being overwhelmed by metabolic demands imposed by mTORC1/oxidative phosphorylation upregulation, a marker of maladaptation. The RV transcriptomic signature (Figure 6D) is fundamentally different from that of lung. RV metabolic upregulation co-exists with downregulation of structural integrity programs, including apical junction assembly, Wnt/β-catenin survival signaling, and hedgehog-mediated repair capacity, indicating simultaneous progression toward maladaptive remodeling.

Where lung shows inflammatory destruction, the RV shows transcriptomic portrait of metabolic reprogramming under stress and early, compensated RV pressure overload transitioning toward maladaptive remodeling driven not by inflammation but by hemodynamic stress from upstream pulmonary vascular stiffening. The RV is working harder, burning fuel differently, losing structural integrity, and beginning to remodel but has not yet failed.

Collectively, these findings define a pathophysiological framework in which a single silica exposure initiates a non-resolving pulmonary inflammatory response that propagates along the lung-heart axis, producing subclinical but measurable dysfunction spanning the alveolar-capillary membrane, the pulmonary vasculature, and the right ventricle, establishing a mechanistic basis for the excess cardiopulmonary morbidity observed in silica-exposed populations.

### Spectral-spatial niche discovery

Although early effects of silicosis manifest at multiple points along the pulmonary circuit, the relative power of these physiologic biomarkers may point to tangible outcomes in our understanding and diagnosis of this pathology. To fully leverage this rich histological dataset, we employed a spectral-spatial 3D wavelet-guided clustering algorithm to uncover geodesic niches that can be ascribed to functional tissue units (Figure 7A). This affords the unified tracking of morphological changes and structures across samples and identifies changes in cell type abundances locally. To capture the cellular relevant cell identities, a supervised vision transformer (CellViT)--a deep encoder-decoder neural network for nuclear segmentation—segmented the cells in the Movat’s pentachrome images (Figure 7B). Although this model was not trained on mouse lung tissue, it retains multiclass probabilities for cell types such as connective, inflammatory, and fibroblast cells. Fibroblast weights were manually tuned, and a sequential structuring element opening-closing operation and subsequent constrained-donut expansion was developed as described in the methods. At the cell level, morphological features such as area, eccentricity, solidity, constrained dilation, and local anisotropy drove the cell type annotations. These segmented cells were embedded in a UMAP using their morphology, colors, and local textures (Figure 7C). There is clear separation between tissue type, but sex, cell type, and treatment condition dimensions were more mixed, yet somewhat separable visually. Although the interactome between fibroblasts and inflammatory cells (a broad mix of various immune lineages) did not materially change between silica and control murine lungs, their interactions with themselves and other cell types in the lumen changed modestly (Figure 7D). Given the abundance of fibroblasts increases in the silica mice, it is no surprise that their distribution is more diffuse and mobilized, as their interaction with lumen increases (Figure 7E).

**Figure 7.**
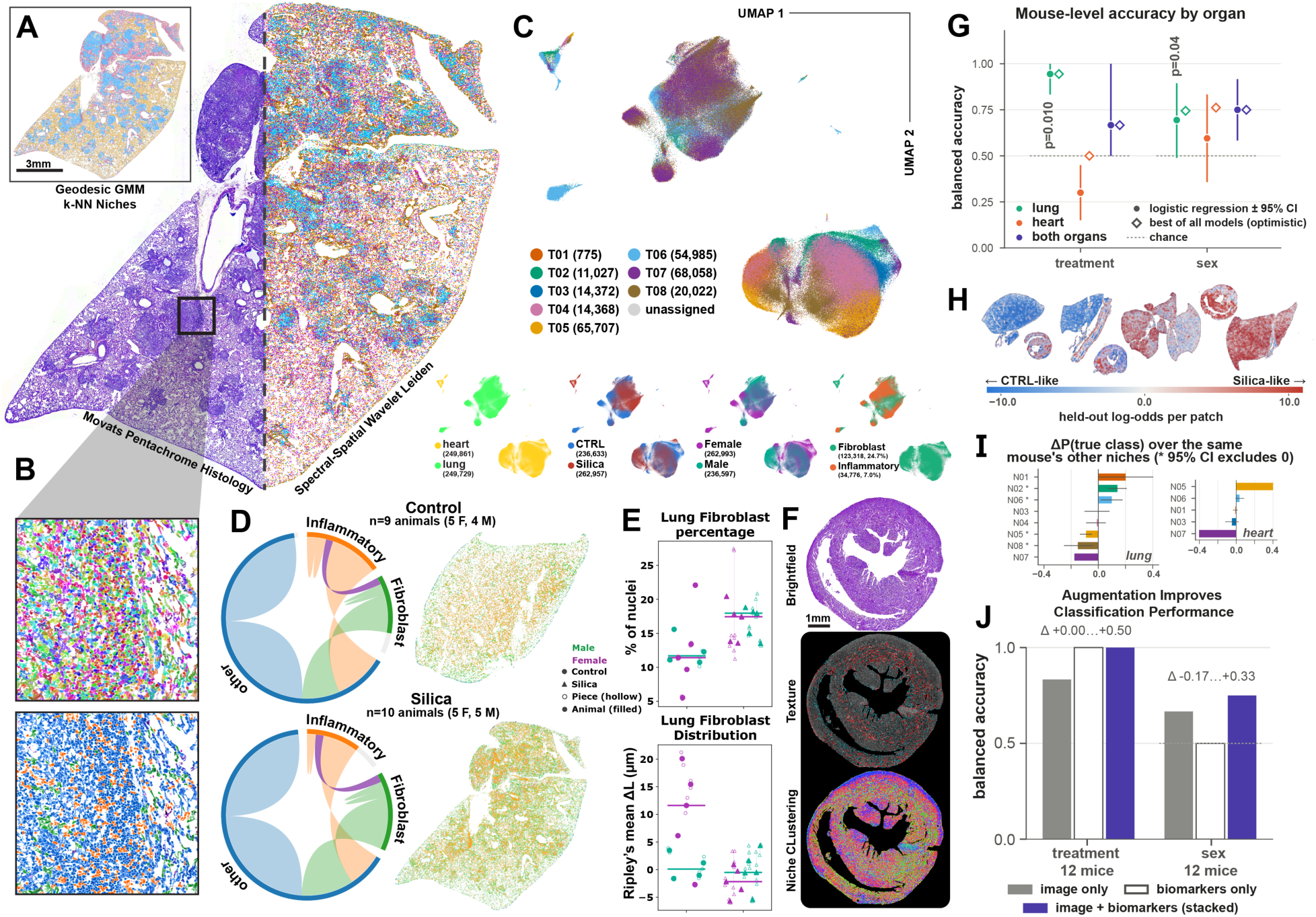
Cell-and tissue-level histology analysis highlights spatial phenotypes of silica-exposed heart and lung tissue. Unsupervised clustering of geodesic tissue niches in Movat’s pentachrome sections (shown: representative silica-exposed male murine lung) derived from spatial-spectral (3D wavelet decomposition, k-means, k=8) using a hierarchical kNN GMM (Panel A). Histology images at 40x resolution were segmented using CellViT and tuned with human-in-the-loop for fibroblast annotations (Panel B). UMAP of 500,000 randomly subsampled cells, colored by shared niche and texture class, with various dimensions highlighted, such as organ, treatment, sex, cell type (Panel C). Cell-cell interactome from Delaunay triangulation (edges ≤ 50 µm). Contact enrichment is shown relative to label-permutation nulls (Panel D). Accumulation and spatial distribution of fibroblasts and inflammatory cells altered after silica exposure. Ripley’s ΔL(r) (r = 10–200 µm) is shown relative to permutation nulls (Panel E - lung). Representative image of silica-exposed murine heart tissue shows cardiac fibrosis gradient ridges. Highlighted pixels are classified as radial or circumferential relative to the ridge (Panel F). Treatment classification from DenseNet-121 features of 224 × 224 µm tiles. Models were trained on lung, heart or both tissues and evaluated by leave-one-mouse-out cross-validation (Panel G). Bootstrapped locations of high-attention patches (class activation maps) and the change in niche share relative to the tissue prior (shown: treatment condition) (Panel H). Niche contributions to classification. The y-axis is the change in predicted probability of the true class, ΔP (true class), for tiles in each niche compared with tiles from the same mouse’s other niches. Values are means with mouse-level bootstrap 95% CIs; asterisks mark niches whose CI excludes 0 (Panel I). Fusion of image-based predictions with physiological biomarkers, each entered as one value per test, show improved classification accuracy. Scale bars are as marked. Error bars show bootstrap 95% Cis (Panel J).

At the tissue level, our spectral-spatial niche discovery also uncovers broad structural gradients as an intermediate consequence. For instance, the heart tissues exhibit distinct gradient separation between radial and circumferential clefts and striations (Figure 7F). When raw histology images were fed to a DenseNet neural network, the lung tissues performed better than the heart tissues at classification accuracy of both silica treatment and gender, confirming the prior observations on sex-dependent physiological responses (Figure 7G). When lung and heart tissues were analyzed in tandem, the accuracy for sex classification improved beyond the lung alone. When examining the locations of the bootstrapped patches fed to the DenseNet, the regions that contributed to a classification are revealed. Interestingly, certain features of the tissues, such as near the ECM of the lung and near the RVFW contained a high concentration of silica-leaning patches. The niches contained in those patches also exhibited a higher change in posterior assignment probability when held out and warrant further exploration into the efficiency of those niches and their diagnostic power (Figure 7I). Augmenting the image data with physiologic biomarker measurements detailed in the previous sections, the accuracy also improves (Figure 7J).

## DISCUSSION

Using a single endotracheal crystalline silica exposure in C57BL/6J mice, we performed integrated cardiopulmonary and molecular phenotyping using bulk RNA-seq of the lung and RV at 28 days. We hypothesized that a single high-burden endotracheal crystalline silica exposure produces persistent lung injury and inflammation that would lead to pulmonary vascular and RV compensatory remodeling. We found that despite apparent recovery of body weight and voluntary exercise capacity, a high-burden of silica in the airways of the lungs initiates a lung → pulmonary artery → right ventricle (lung-PA-RV) cascade characterized by monocytic-predominant nodular infiltrates and alveolar-capillary dysfunction thereby elevating pulmonary circulatory pressures. Immune-mediated fibroblast This drives immune-mediated fibroblast remodeling of the distal pulmonary vascular remodeling and subsequent proximal pulmonary vascular stiffening. Pulmonary vascular remodeling increases RV afterload and initiates RV remodeling/metabolic reprogramming, with sex-dependent cardiopulmonary phenotypes.

### A single endotracheal silica causes insidious cardiopulmonary mechanical abnormalities initiated by lung injury, leading to pulmonary vascular and cardiac maladaptive changes: a lung-PA-RV axis of maladaptive remodeling

Our data shows a clear temporal cascade initiated by endotracheal silica-induced lung injury and manifests 28 days later as functional pulmonary artery stiffening and right ventricular compensation. A subclinical phenotype is noted, demonstrating the insidious nature of the cardiopulmonary remodeling process. There is a disassociation between apparently preserved cardiopulmonary performance and objective physiological injury one month after the insult. This is like human phenotypes of workers and veterans exposed to silica and other substances who experience a latency period during which they appear normal in clinic and on routine tests as their chronic lung injury-induced disease progresses. In these and similar cases, homeostatic inflammatory responses initially serve compensatory functions that preserve tissue integrity and organ/system function, e.g., cardiopulmonary exercise function; however, persistent activation can transition to maladaptive inflammation that promotes tissue dysfunction and disease progression (*39*).

We found multiple independent readouts consistent with ongoing lung injury 28 days after the initial insult. Together, these argue that the initial insult is pulmonary in nature, and measurable, structural, and viscoelastic abnormalities are left behind throughout the cardiopulmonary system. While many other investigators have demonstrated the profound effects that these exposures can have on isolated organs or tissue, our study is amongst the first to rigorously support the theory of a lung injury propagating to proximal pulmonary vascular remodeling, to RV remodeling (28).

Silica in the airways induces lung injury and inflammation that leads to alveolar-capillary endothelial / alveolar epithelial type II barrier leak which impairs gas exchange. Together, these lead to remodeling of the distal pulmonary vasculature in the form of increased muscularization and collagen deposition consistent with hypoxic constriction, which drives the increased pulmonary circulatory pressures. This pressure increase drives the elevation of circumferential stress in the arterial wall. While primarily studied in systemic circulation, there is a strong body of literature supporting tight regulation of circumferential wall stress as a homeostatic setpoint, and sustained elevation in transmural pressure drives growth and remodeling that return wall stress toward that baseline (*40*). Such regulation would explain why the pulmonary arteries of silica-exposed mice exhibited lower circumferential stress than controls when evaluated at identical pressures. The remodeling observed here is predominantly geometric: wall thickness trended upward and the wall-to-lumen ratio increased, offloading wall stress as expected from its inverse relationship to Laplacian wall stress. That this geometric remodeling is already apparent 28 days post-exposure raises the possibility that a material-stiffening component emerges at later timepoints, one that would compound the pulsatile load imposed on the right ventricle. This motivates longitudinal studies spanning longer exposure durations. At 28 days, however, the altered proximal PA stiffness and pulse wave velocity appear attributable almost entirely to the elevated operating point rather than to intrinsic material remodeling of the wall.

The reduced distensibility of the proximal pulmonary artery at *in vivo* pressures limits its capacity to participate in the Windkessel effect (mechanical elastic buffering). This results in a greater fraction of pulsatile energy being transmitted into the distal pulmonary microvasculature, potentially further impairing gas exchange while simultaneously increasing the pulsatile component of RV afterload. Downstream right ventricular cardiac effects in both structure (free wall thickness) and remote molecular remodeling were observed as a result.

The multiple levels of concordance between physiology, biomechanics, hemodynamics and transcriptomics support a lung-PA-RV axis of injury. This not only demonstrates the chain of events that may lead to latent cardiopulmonary impairment, but it also identifies structural, functional, and molecular targets for earlier identification, better prognostication, and potential therapeutic interventions during this latent period. Our observations of changes along the lung-PA-RV axis of injury may reflect compensatory changes in the cardiopulmonary system that, if they persist, lead to maladaptive remodeling.

### Sex as a biological variable plays a role in lung tissue responses to injury

At 28 days after the same single endotracheal silica exposure, both sexes show clear evidence of persistent lung injury (reduced DFCO, increased BAL cellularity with elevated BAL protein, foamy macrophages, and ultrastructural evidence of barrier disruption). This establishes a shared response to lung airway inhalation injury. However, the downstream pulmonary phenotype diverges by sex: females exhibit a parenchymal-dominant mechanical signature (↑hysteresis, ↓tissue elastance, ↑ hysteresivity, ↑inspiratory capacity), whereas males exhibit an airway-dominant signature (↑ Newtonian/airway resistance). Importantly, the “female-type” parenchymal parameters are not significantly altered in males, and the “male-type” airway resistance is not significantly altered in females, indicating qualitatively different compartment-level responses rather than a simple difference in severity.

These functional differences are reinforced anatomically and are not readily explained by body size. The multinodular, monocyte-predominant peribronchiolar infiltrative process is present overall, but males have a higher proportion of airways filled with infiltrative debris, consistent with their increased airway resistance. Because males are typically heavier and larger, particle deposition is plausibly confounded by airway size because airway size increases as body mass and size increase based on normative aging and sex (*30*). However, allometric analysis indicates the observed sex divergence is independent of body mass, supporting sex-linked biology as the driver of the sex-based differences we observe. Finally, sex also appears to modify downstream cardiopulmonary remodeling: silica exposure increases RV pressure, and this associates with increased RV free wall thickness in females, suggesting sex-dependent adaptation to the afterload imposed by lung injury–associated pulmonary vascular remodeling.

### Silica-induced lung injury elicits an inflammatory response in the lung, while the right ventricle undergoes a shift toward metabolic reprogramming

In the lung, co-enrichment of interferon alpha, interferon gamma, IL-6/JAK/STAT3, and inflammatory response pathways alongside downregulation of Kras and myogenesis programs at 28 days post-silica exposure defines a transcriptomic state of persistent, maladaptive immune-driven inflammation. This pattern is consistent with a maladaptive inflammatory-fibrotic transition. Ongoing immune-mediated tissue injury is not offset by compensatory epithelial or structural regeneration. The prominence of IL-6/STAT3 and interferon signaling, which was previously found to activate perivascular macrophages and adventitial fibroblast (*22, 41*), provides a plausible mechanistic link between the lung parenchymal inflammatory response and pulmonary vascular remodeling due to increased pulmonary circulatory pressures. This suggests that perivascular macrophage-driven vascular maladaptive remodeling may be a downstream consequence of non-resolving silica-induced lung inflammation, supported by our analyses that suggests inflammatory signaling of fibroblast in the lung associated with silica exposure.

The RV GSEA results suggest the cytoprotective capacity of RV cardiomyocytes is being overwhelmed by metabolic demands imposed by mTORC1/oxidative phosphorylation upregulation, a marker of maladaptation. The RV transcriptomic signature is fundamentally different from that of lung. RV metabolic upregulation co-exists with downregulation of structural integrity programs, including apical junction assembly, Wnt/β-catenin survival signaling, and hedgehog-mediated repair capacity, indicating simultaneous progression toward maladaptive remodeling. Silica-specific remodeling associates with localized remodeling of the right ventricle free wall and interventricular septum, suggesting that cardiac remodeling due to silica exposure is driven by remodeling of the pulmonary vasculature rather than intrinsic cardiac dysfunction.

Where lung shows inflammatory injury, the RV shows transcriptomic signatures of metabolic reprogramming under stress; early, compensated RV pressure overload transitioning toward maladaptive remodeling driven not by inflammation but by hemodynamic stress from upstream pulmonary vascular stiffening. The RV is working harder, burning fuel differently, losing structural integrity, and beginning to remodel but has not yet failed.

### Limitations

Key limitations of this study are that crystalline silica alone may evoke a comparatively acute response compared to repeated, lower-dose exposures (*42*). Exposure other than silica may also elicit different effects due to the complex mixture of deployment-relevant dusts. Therefore, translation of these findings to deployment-related respiratory diseases in veterans are limited silica dust only. These findings raise several questions for future work, including defining the time course linking initial lung injury to subsequent pulmonary vascular and RV remodeling, determining whether the observed “insidious” mechanical abnormalities persist or resolve, and extending to later time points to test whether the RV transcriptomic signature currently suggestive of compensation rather than the Warburg-like profile associated with RV failure (*43*). It is unclear if these early compensations eventually transition to decompensation or maladaptation and on what timeline. Additional priorities are to evaluate whether the sex-dependent phenotypes and other readouts translate to human cohorts, and to determine whether the lung-PA-RV axis contains compartment-, organ-, or cell-specific therapeutic or mechanistic targets.

## Author Contributions

**Conceptualization:** MSG, JD, ZMH, and EPM

Methodology: MSG, JD, AAF, RC, PHG, AAS, PKL, XL, NG, XY, ZMH, and EPM

Investigation: MSG, JD, AAF, RC, PHG, AR, AN, AAS, YS, KH, PKL, XL, ZMH, and EPM

Analysis: MSG, JD, AAF, RC, PHG, AR, AN, AAS, TS, SLB, BH, YS, PKL, XL, NG, XY, JMS, SMM, RF, ZMH, and EPM

Visualization: MSG, JD, AAF, PHG, AR, AN, AAS, XL, SMM, ZMH, and EPM

Writing: MSG, JD, AAF, SMM, ZMH, and EPM drafted the manuscript; MSG, JD, AAF, RC, PHG, AR, AN, AAS, TS, YS, SLB, BH, YS, KH, CB, PKL, JG, XL, NG, JLK, XY, JMS, SMM, RF, ZMH, and EPM approved final version of the manuscript.

## Disclosures/Competing interests

EPM reports consulting for Biomedical Consultants, PLLC, outside the submitted work.

## Funding

T32HL007778 to MSG. The research reported/outlined here was supported by the Department of Veterans Affairs, Veterans Health Administration, VISN 1 Career Development Award to Dr. Zachary Harris.

## Notes

https://github.com/jfd-lab/silica_bulkRNAseq

