## Supplemental methods and figures for "Crystalline silica exposure causes latent biomechanical abnormalities along the cardiopulmonary axis"

**Supplemental material for Crystalline silica exposure causes latent biomechanical abnormalities along the cardiopulmonary axis**

Meredith S. Gwin^1^*, Jack Di Palo^1^*, Anthony A. Fung^2^*, Rira Choi^2^, Pete H. Gueldner^2^, Alexandria Regan^1^, Aishani Nagamalla^1^, Asad A. Syed^1^, Theodore Sandow^1^, Ying Sun^1^, Sydney L. Bluestein^1^, Buqu Hu^1^, Yosep Shin^1^, Keshav Harpavat^1^, Clemente Britto^1^, P. Kent Langston^3^, Jose Gomez^1^, Xinran Liu^4^, Nicole Guerrera^5^, Jonathan L. Koff^1^, Xiting Yan^1,6^, Jason M. Szafron^7^, Susan M. Majka^8^, Rong Fan^2^, Zachary M. Harris^1,9 †^, and Edward P. Manning^1,9†^

*****, † indicates equal contribution

1. Section of Pulmonary, Critical Care and Sleep Medicine, Yale School of Medicine, Yale University, New Haven, CT, USA

2. Department of Biomedical Engineering, Yale University, New Haven, CT, USA

3. Department of Pathology, Yale School of Medicine, Yale University, New Haven, CT USA

4. Center for Cellular and Molecular Imaging Electron Microscopy Facility, Department of Neuroscience, Yale School of Medicine, Yale University, New Haven, CT USA

5. Department of Medicine (Cardiovascular Medicine), Yale Translational Research Imaging Center, Yale School of Medicine, Yale University, New Haven, CT, USA

6. Department of Biostatistics, Yale School of Public Health, New Haven, CT, USA

7. Department of Biomedical Engineering, Carnegie Mellon University, Pittsburgh, PA, USA

8. Division of Pulmonary, Critical Care & Sleep Medicine, Department of Medicine, National Jewish Health, Denver, CO, USA, Gates Center for Regenerative Medicine and Stem Cell Biology, University of Colorado, Aurora, CO, USA

9. VA Connecticut Healthcare System, West Haven, CT, USA

**Supplemental Methods**

**Mice**

Animal experiments were designed, conducted, and reported in accordance with the ARRIVE 2.0 guidelines to promote rigor, transparency, and reproducibility in preclinical research (*1*). It was approved by the Yale University Institutional Animal Care and Use Committee. Female and male C57BL/6J WT mice aged 13-15 weeks old (to model a young-adult life stage, which is most analogous to the typical age range of U.S. military service members during deployment and occupational inhalation exposures) were housed in an antigen‐free and virus‐free animal care facility under a 12:12 light:dark cycle. Mice were fed a standard rodent chow and had free access to food and water. Mice were randomized to receive either silica exposure or vehicle control, and experiments were repeated to confirm reproducibility. To minimize potential hormonal effects in females, exposures were performed at varying, randomly assigned points across the estrous cycle. Investigators were blinded to treatment group by using coded animal identifiers, and group assignments were not revealed until all analyses were completed.

**Silica**

Prior to use silica was heated to 500° F for at least 2 hours and then resuspended in sterile 0.9% saline under aseptic conditions. Before each experiment the silica was vortexed well, aliquoted into 2mL vials, and homogenized using an ultrasonic homogenizer for 1-2 minutes.

**Exposure**

Silica-exposed mice were endotracheally administered a single dose of crystalline silica (0.4 g/kg of bodyweight) suspended in a 0.9% saline solution vehicle. Control mice were administered only the vehicle. The total volume of solution administered was 3.33 μL/g of bodyweight. The dose was selected as a high-burden exposure model intended to mimic the cumulative pulmonary deposition resulting from repeated exposure to silica-containing desert dust encountered during prolonged deployment in Southwest Asia or a bolus experienced during severe sandstorms. Dusts from Iraq and Afghanistan contain quartz and other silica-bearing minerals, and airborne particulate concentrations can rise dramatically during severe dust storms (*2-4*). The bolus intratracheal administration represents a compressed model of chronic cumulative lung burden over a year-long deployment when the average 2.5 micron-sized airborne particulate concentration is ~100 ug/m^3^ (*5*) or a single environmental exposure event such as a severe sandstorm when silica particulate matter concentration exceeds 10,000 ug/m^3^ (*2-4*). This dose has been shown to cause pulmonary hypertension in C57BL6 mice (*6*).

Endotracheal administration was confirmed by using a sialendoscopy scope (PolyDiagnost, Hallbergmoos, Germany) placed within the cannula and advancing the scope-cannula system until the carina of the subject could be visualized, after which, the scope was removed, leaving just the cannula in place and allowing for bolus administration.

**Echocardiography**

Noninvasive investigation of cardiac function was performed in mice using transthoracic echocardiograms under light anesthesia (1.5% isoflurane) while maintaining physiological temperature (*7*). Chest fur was removed using Nair depilatory cream. Body temperature and heart rate were monitored throughout the procedure. At the end of the procedure mice were returned to their home cages and allowed to recover. Standardized cardiac views were obtained with a high‐resolution ultrasound system (Vevo 2100; VisualSonics, Toronto, ON, Canada) equipped with an ultrahigh frequency (40 MHz) linear array transducer. B‐mode two‐dimensional (2D) images of the pulmonary artery were obtained from a parasternal short‐axis view at the level of the aortic valve. In addition, M‐mode and tissue Doppler imaging (TDI) of the lateral tricuspid annulus were obtained from a modified apical four‐chamber view. The pulmonic valve was imaged at the level of the leaflet tips with pulsed wave Doppler. Tricuspid annular plane systolic excursion (TAPSE), RV systolic myocardial velocity (s'), pulmonary artery acceleration time (PAT), and pulmonary ejection time (PET) were measured offline using Vevo Lab software (version 3.2.6, VisualSonics).

**Diffusion Factor for Carbon Monoxide Measurement**

Diffusion Fraction for Carbon Monoxide of the lung (DFCO) measurements were made using a gas chromatographer (Inficon, Micro GC Fusion) (*8*). Mice were anesthetized with urethane (intraperitoneal injection, 1.5–2.0 g/kg), and the depth of anesthesia was assessed as the lack of response to a toe pinch, with supplemental injections given as needed. Once adequately anesthetized, the mouse was placed supine, and the trachea was cannulated with a 20-gauge tracheostomy tube. The tube was inserted through a small ventral incision made in the rostral-most part of the trachea and advanced 3 mm caudal to the incision. The tube was then held securely in place with a suture tied around the trachea. 600 µL of 0.5% carbon monoxide, 0.5% neon molar fraction air-balanced gas was pushed into the mouse lungs, allowed to dwell for 9 seconds, then 600 µL was pulled from the lungs back into the syringe which was then disconnected from the tracheal tube. 1400 µL of room air was pulled into the syringe and a finger was quickly placed over the opening. The syringe was then connected to the gas chromatographer, and the syringe was depressed. The diffusion factor for carbon monoxide was calculated as 1 – (CO_9_ / CO_c_) / (Ne_9_ / Ne_c_), where the c and 9 subscripts refer to molar concentrations of the gases injected and the gases removed after a 9 second breath hold time, respectively. DFCO is a dimensionless variable, which varies between 0 and 1, with 1 reflecting complete uptake of all CO, and 0 reflecting no uptake of CO.

**Pulmonary Function Testing**

Following DFCO measurements, the surgical plane of anesthesia was reconfirmed as lack of response to toe pinch, with supplemental injections of Urethane given as needed. Following confirmation, the tracheostomy cannula was connected to the FlexiVent system (SCIREQ, Montreal, QC, Canada) for a pulmonary function test (PFT). Prior to beginning the PFT, pancuronium bromide (1 mg/kg), a neuromuscular blocker, was administered via intraperitoneal injection to render the mouse’s breathing passive. Once passive breathing had been established, the PFT maneuvers began using a forced-oscillation technique (FOT). A positive end-expiratory pressure of 3 cmH_2_O was maintained throughout the PFT. The pulmonary function test revealed parameters for inspiratory capacity (IC), static compliance (Cst), dynamic compliance (Crs), elastance (Ers), tissue elastance (H), tissue damping (G), and Newtonian resistance (Rn). Additionally, pressure-volume curves were generated, from which the area between the loading and unloading curves could be measured as hysteresis. PFT parameters were extracted from the raw data with SCIREQ software (Flexiware v.7.6, Service Pack 6).

**Bronchoalveolar Lavage (BAL) Fluid Collection and Analysis**

After pulmonary function testing mice were placed supine with limbs splayed and taped down. A midline cut was made in the abdomen, and the intestines and liver were moved such that the diaphragm could be visualized. The diaphragm was cut from left side to right making sure not to cut the lungs or heart. Two cuts were made on either side of the ribcage up to below the clavicle. Needle holders were used to fold the ribcage upward and out of the field of view of the heart/lungs. The mice were exsanguinated and perfused with chilled Hanks buffered saline solution (HBSS, ThermoFisher 14025). Then, the rest of the chest was removed to expose the trachea. Using the tracheotomy that was previously done, a 1mL luer-lock syringe containing 800 μL of ice-cold Dulbecco’s phosphate-buffered saline (DPBS) (Gibco 14190-144) was attached to the canula and the fluid instilled slowly. DPBS was removed and collected into a sterile labeled 2mL microfuge tube. This was repeated once more for a total of 1.6mL instilled per mouse. The BAL fluid was kept on ice until the end of the procedure then total cell counts were made using an automated COULTER cell counter (Beckman Coulter, Brea, CT, USA) or the ThermoFisher countess (AMQAF1000). BAL was centrifuged at 450 X g for 10 minutes to pellet cells. The supernatant was removed and placed into new sterile microfuge tubes, flash frozen and stored at -80 until use. The cell pellet was then resuspended in 100 μL DPBS and cytospun for histology.

Cytospin slides were stained using Diff-Quik (Dade Behring AG, Düdingen, Switzerland) to allow for evaluation of BAL cells and identification of macrophages, lymphocytes, neutrophils, eosinophils, and basophils (*9*). Cell differential analysis was completed by counting cells contained in three 20x high-powered fields per sample. The individual counting cells was blinded to the experimental conditions of each sample. An average of 840 total cells were counted per sample. Absolute cell count for specific cell types (e.g., macrophages, lymphocytes, etc.) was determined by multiplying the cell differential percentage for that cell type with the BAL total cell count obtained as described in the section above. BAL total protein, a marker of lung injury and loss of alveolar-capillary barrier integrity (*10*), was measured using the bicinchoninic acid (BCA) assay (ThermoFisher, product #23225).

**Proximal Pulmonary Artery Biomechanical Measurement and Analysis**

Following collection of the BAL fluid, the heart and lungs were removed from the thoracic cavity *en bloc* and lateral pieces of each of the lobes of the right lung and RV were removed, flash-frozen, and stored at -80 °C until later use. The rest of the heart and lungs were stored temporarily in HBSS at 4 °C. Afterwards, fat and perivascular tissue were removed from the heart-lung block to reveal the pulmonary artery, attached to the heart and branching to both lungs. The left pulmonary artery branch and ligamentum arteriosum were tied off with suture. The pulmonary artery was then cannulated through the main and right branches with glass micropipettes and mounted on a custom biaxial computer-controlled testing device (*11*). The mechanical tests, consisting of two separate inflation protocols at fixed axial lengths (1.05, 1.00, 0.95) and axial extensions at fixed pressures (5, 15, 25, 40 mmHg) have been described previously in other literature (*12*). Vessel diameter was automatically tracked by a camera, and vessel extension was affected with micro stepper motors. Pressure and force transducers were used to measure intraluminal pressure and resistance to axial extension/compression, respectively. During the tests, the vessel was submerged in room temperature HBSS, whose low calcium concentration renders smooth muscle cells passive (*13*). The raw data was fit to the four-fiber-family constitutive model for arterial walls (*14, 15*), providing eight material parameters from which clinically relevant mechanical properties of the artery may be derived. The methods of parameter estimation has been previously described in the literature (*12*).

**Bulk RNA sequencing**

RNA was extracted from the flash-frozen samples of lung and RV, stored at –80 °C using miRNeasy Micro kits (Qiagen, 217084) following manufacturer’s protocol. Purified total RNA was submitted to Plasmidsaurus (Eugene, OR) for 3′-end counting RNA sequencing. RNA concentration was assessed by a fluorescence-based plate-reader assay and normalized across samples prior to library preparation. Libraries were prepared by reverse transcription of mRNA using a poly(dT)VN primer, followed by second-strand synthesis, tagmentation with a transposase to simultaneously fragment the cDNA and add sequencing adapters, library indexing, and amplification. Unique molecular identifiers (UMIs) were incorporated during library construction and unique dual indices (UDIs) were used to prevent index hopping. Libraries were sequenced on Illumina NovaSeq instruments in single-end, stranded configuration, with reads oriented in the same sense as the input mRNA and sequencing directed toward the transcript 3′ end. Reads are approximately 90 nt in length and align within roughly the final 400 nt of each transcript. Base calling and demultiplexing were performed by the vendor using BCL Convert (v4.3.6) and fqtk (v0.3.1), and raw FASTQ files were retrieved for independent processing.

Read quality was assessed with FastQC (v0.12.1) and aggregated across samples with MultiQC. Adapter sequence, poly-A read-through, and poly-G artifacts were removed using Cutadapt (v5.1) in three sequential passes: (i) trimming of the Illumina universal adapter (AGATCGGAAGAG) with quality trimming (-q 15); (ii) poly-A tail removal (--poly-a); and (iii) NextSeq/NovaSeq poly-G trimming. In each pass, reads shorter than 30 nt after trimming and reads containing more than two ambiguous bases were discarded. Read quality was re-assessed after trimming. Trimmed reads were aligned to the mouse reference genome GRCm39 using the GENCODE release M39 primary-assembly annotation. A STAR (v2.7.11b) genome index was generated to the maximum observed read length (94 nt). Alignment was performed with STAR in two-pass mode. Only samples with an alignment rate greater than 65% were kept and analyzed.

Aligned reads were assigned to genes using featureCounts (Rsubread v2.1.1) against the GENCODE M39 annotation. Count matrices were assessed for library size, number of genes detected at ≥10 reads, proportion of reads in protein-coding genes, mitochondrial read fraction, and the fraction of reads attributable to the single most abundant gene.

Right-ventricular tissue was subject to variable residual blood content, which can dominate count distributions. The proportion of counts derived from hemoglobin genes (Hbb-bs, Hbb-bt, Hba-a1, Hba-a2, Hbb-y, Hbb-bh1) was therefore computed for each RV sample and included as a continuous covariate in the RV differential-expression model.

Differential expression between crystalline silica-exposed and control animals was assessed separately for each tissue using DESeq2. Genes were retained if they reached at least 10 counts in at least three samples, corresponding to the size of the smallest experimental group. Both sexes were included, with design formulas ~ Sex + Exposure for lung and ~ Sex + pct_hb + Exposure for RV, using control as the reference exposure level. Genes were considered differentially expressed at a Benjamini–Hochberg adjusted p-value < 0.05 and |log2 fold change| ≥ 1.5.

Gene set enrichment analysis (GSEA) was performed with clusterProfiler using Mus musculus Hallmark gene sets. Genes were ranked by log2 fold change across the full detected transcriptome, and enrichment significance was assessed at a Benjamini–Hochberg adjusted p-value < 0.05.

Processing was performed on the Yale Center for Research Computing Bouchet cluster. Software versions were: FastQC 0.12.1, MultiQC 1.28, Cutadapt 5.1, STAR 2.7.11b, Rsubread 2.1.1, R 4.4.2, Bioconductor 3.20, DESeq2 1.24.0, clusterProfiler 4.10.0, and msigdbr 25.1.1. Analysis code is available at <https://github.com/jfd-lab/silica_bulkRNAseq>; raw sequencing data have been deposited in the NCBI Gene Expression Omnibus (GEO) under the accession number GSE344989.

**Pathological biomarker classification and analysis**

**Tissue niches.** Texture classes of Movat's pentachrome histology came from a Gabor filter bank (wavelength 2–6 µm, 30° steps) clustered by k-means (k = 8). Spectral-spatial classes came from a 3D wavelet decomposition (three spatial levels and one spectral level) clustered by k-means (k = 8). Geodesic niches representing functional tissue unit (FTU) level segmentation were built by diffusing cell features over a 12-nearest-neighbour graph at 8, 20 and 45 µm, then fitting a Gaussian mixture (8 niches). All three label sets were harmonized across pieces with a shared codebook and re-clustered per organ into 8 shared types at the organ level.

**Cell segmentation.** CellViT (SAM-H encoder, PanNuke head) segmented nuclei directly from the 40x Movat's pentachrome images. Cell bodies came from opening-closing a cytoplasm mask (1 and 2 µm), then a constrained-donut watershed expansion within that mask (≤ 10 µm). Inflammatory calls required P ≥ 0.99. The fibroblast threshold was tuned with a human in the loop: overlays were compared repeatedly with raw-image ROIs until the calls matched morphology (final P ≥ 0.9).

**Embedding.** 500,000 randomly subsampled cells (without replacement) were embedded by UMAP (30 neighbors, min_dist 0.1) and clustered by Leiden (resolution 1.0).

**Spatial statistics.** Interactomes came from Delaunay triangulation of cell centroids (edges ≤ 50 µm), with contact frequencies tested against label permutations. Ripley's ΔL(r) = L_obs − L_null was computed for fibroblasts and inflammatory cells (r = 10–200 µm; 39 permutations; ≤ 5,000 subsampled cells per piece). Group differences used 4,999 animal-label permutations. In the heart, ridges were taken from the gradient magnitude of smoothed fibroblast density, and fibers were classed as radial or circumferential relative to the ridge normal.

**Classification.** Tiles of 224 × 224 px at 1.0 µm/px (≥ 50% tissue, ≤ 4,000 per piece) were Reinhard-normalized and encoded by DenseNet-121 (ImageNet weights, fine-tuned for 4 epochs on 200 tiles per slide). Features were reduced by PCA to 128 dimensions and aggregated per mouse. Analyses were pooled by tissue type (lung, heart, both). Ten classifiers were evaluated by leave-one-mouse-out cross-validation, with 100 label permutations and 2,000 mouse-bootstrap CIs. They included logistic regression, SVMs, MLP, random forest, gradient boosting and a color-only control. Class activation maps (7 × 7 grid, ~32 µm cells) were projected onto niches. A niche was significant when its CAM-weighted share minus the tissue prior had a mouse-bootstrapped 95% CI excluding zero.

**Fusion.** Each physiological test or measurement contributed one biomarker, such as RVFW thickness, DFCO, etc. Body mass was summarized as the mean % of baseline across days, and multi-measure tests as their first principal component. Biomarkers were ranked by permutation importance and mutual information, then combined with image probabilities by late fusion and stacking under leave-one-mouse-out cross-validation.

**Supplemental Figure S1**

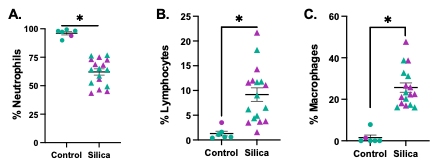

**Supplemental Figure S1:** BAL cell differential demonstrated significantly decreased percentage of macrophages and significantly increased percentages of lymphocytes and neutrophils compared to saline controls. * = *P* < .05. n = 6-17 per group.

**Supplemental Figure S2**

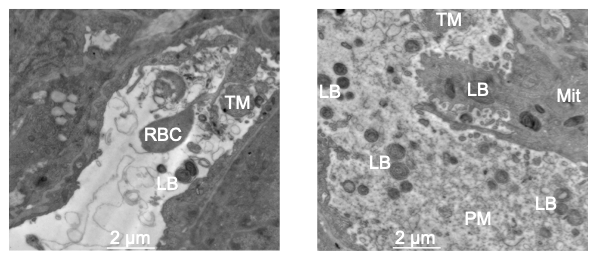

**Supplemental Figure S2:** Representative transmission electron microscopy images of lungs from mice treated with silica show red bloods, non-degraded surfactant, lamellar bodies, tubular myelin and proteinaceous material in the airways, indicative of a disrupted alveolar-capillary membrane. RBC = red blood cell, TM = tubular myelin, LB = lamellar body, Mit = mitochondria, and PM = particulate matter.

**Supplemental Figure S3**

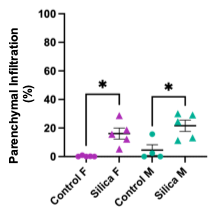

**Supplemental Figure S3.** Mice exposed to silica had a significantly increased percentage of peribronchial infiltrative processes of the parenchyma (*P* = .0146 between females and *P* = .0167 between males) with male mice having a greater percentage (21.6% vs females with 16.1%, *P* = .3464). n = 4-5.

**Supplemental Figure S4**

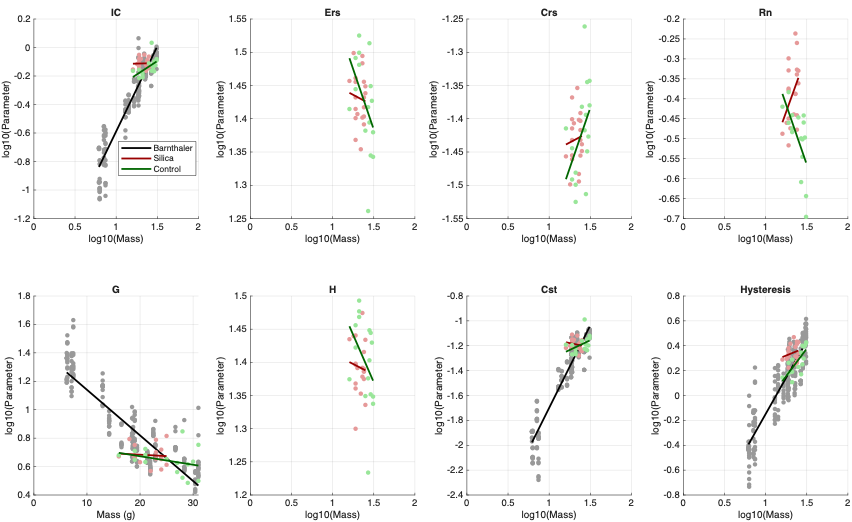

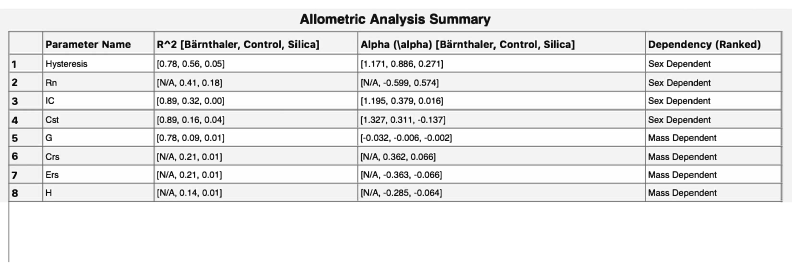

**Supplemental Figure S4: Silica exposure disrupts normal allometric scaling, demonstrating that differences in lung function between males and females may not be explained by body size alone.** The top panel shows log-log allometric regressions comparing log10(body mass) against log10(parameter) for inspiratory capacity (IC), respiratory system elastance (Ers), respiratory system compliance (Crs), airway resistance (Rn), tissue elastance (H), static compliance (Cst) and hysteresis. Tissue damping (G) is plotted using a semi-log regression (raw mass vs log10(G)). Linear regression models were fit using data from three different cohorts of mice: a historical baseline data set (Bärnthaler et al., black line), healthy saline controls (Control, green line) and silica-exposed mice (Silica, red line)(*16*). Historical data from Bärnthaler et al was only available for IC, G, Cst, Hysteresis. While the healthy cohorts display strong scaling slopes, silica exposure flattens or inverts these allometric relationships across multiple parameters. The table in the bottom panel presents the goodness-of-fit values (R^2^) and scaling exponents (α) of the eight parameters calculated from the linear regressions. For hysteresis, Rn, IC and Cst, the substantial divergence in R^2^ and α values between the silica-exposed mice and healthy cohorts leads us to believe that observed male-female differences in these parameters are driven by biological sex rather than body mass.

**Supplemental Figure S5**

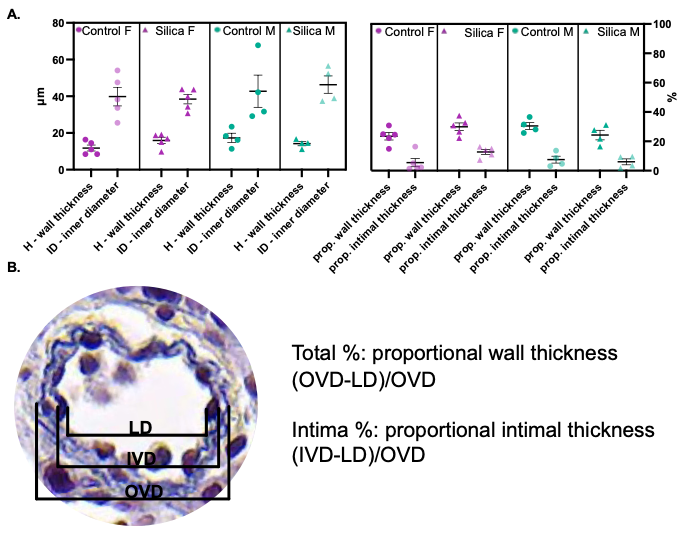

**Supplemental Figure S5: Remodeling of distal intralobar pulmonary arteries (PAs) in silica-exposed mice.** Morphometric analysis of distal PA geometry in vehicle- and silica-exposed mice. Unloaded wall thickness and inner diameter were unchanged; however, trends toward increased intimal thickness were observed in females (Panel A right side, *P* = .0631).

**Supplemental Figure S6**

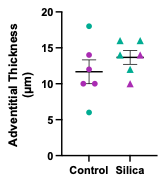
**Supplemental Figure S6: Pulmonary artery adventitia trends towards thickening**

**Supplemental Table S1: Estimated Four-Fiber Family model parameters for the proximal pulmonary artery per group**

|  | **Female Control n = 5** | **Male Control**  **n = 6** | **Female Silica**  **n = 5** | **Male Silica**  **n = 5** |
| --- | --- | --- | --- | --- |
| **Elastic Fibers** c (kPa) | 11.25 ± 1.82 | 6.87 ± 1.40 | 10.15 ± 1.90 | 8.57 ± 0.95 |
| **Axial Collagen** |  |  |  |  |
| c_1_^1^ (kPa) | 6.27 ± 4.04 | 7.30 ± 6.86 | 9.33 ± 3.75 | 11.10 ± 6.02 |
| c_2_^1^ (kPa) | 10.67 ± 3.64 | 6.35 ± 2.55 | 6.79 ± 0.86 | 9.63 ± 2.73 |
| **Circumferential Collagen** |  |  |  |  |
| c_1_^1^ (kPa) | 0.23 ± 0.23 | 0.91 ± 0.46 | 0.24 ± 0.11 | 0.46 ± 0.19 |
| c_2_^1^ (kPa) | 0.60 ± 0.60 | 0.03 ± 0.03 | 0.23 ± 0.23 | 0.00 ± 0.00 |
| **Diagonal Collagen** |  |  |  |  |
| c_1_^3,4^ (kPa) | 9.53 ± 1.75 | 12.30 ± 4.01 | 11.04 ± 2.16 | 12.30 ± 1.44 |
| c_2_^3,4^ (kPa) | 0.96 ± 0.12 | 0.85 ± 0.28 | 1.20 ± 0.22 | 1.07 ± 0.25 |
| α_0_ (deg) | 30.17 ± 2.17 | 33.69 ± 2.78 | 28.65 ± 2.01 | 27.01 ± 1.49 |
| **Error (RMSE)** | 0.08 ± 0.01 | 0.12 ± 0.02 | 0.08 ± 0.01 | 0.08 ± 0.00 |

**Supplemental Table S2: Proximal pulmonary artery geometry and mechanics evaluated at common pressures per group**

|  | **Female Control**  **n = 5** | **Male Control**  **n = 6** | **Female Silica**  **n = 5** | **Male Silica**  **n = 5** |
| --- | --- | --- | --- | --- |
| **Unloaded Dimensions** |  |  |  |  |
| Wall Thickness (μm) | 57.76 ± 3.79 | 57.54 ± 3.79 | 60.60 ± 1.30 | 61.98 ± 1.25 |
| Outer Diameter (μm) | 674.60 ± 46.57 | 675.18 ± 54.70 | 620.60 ± 35.31 | 642.20 ± 21.21 |
| Axial Length (mm) | 2.12 ± 0.12 | 1.95 ± 0.31 | 2.28 ± 0.10 | 2.33 ± 0.16 |
| **Loaded Dimensions** | **P = 15.0 mmHg** | **P = 15.0 mmHg** | **P = 15.0 mmHg** | **P = 15.0 mmHg** |
| Wall Thickness (μm) | 35.44 ± 2.39 | 34.82 ± 2.23 | 39.56 ± 1.61 | 38.50 ± 0.83 |
| Outer Diameter (μm) | 908.67 ± 54.57 | 900.90 ± 42.43 | 816.10 ± 12.60 | 889.52 ± 15.54 |
| Axial Length (mm) | 418.90 ± 26.79 | 415.63 ± 21.49 | 368.49 ± 4.90 | 406.27 ± 7.19 |
| *in vivo* Axial Stretch (λziv) | 1.15 ± 0.03 | 1.18 ± 0.03 | 1.10 ± 0.02 | 1.10 ± 0.02 |
| *in vivo* Circumferential Stretch (λϑiv) | 1.43 ± 0.05 | 1.43 ± 0.09 | 1.41 ± 0.07 | 1.47 ± 0.04 |
| **Cauchy Stresses (kPa)** |  |  |  |  |
| Circumferential (σϑ) | 23.95 ± 1.84 | 24.50 ± 2.33 | 18.73 ± 0.64 | 21.12 ± 0.33 |
| Axial (σz) | 23.92 ± 2.92 | 24.29 ± 3.34 | 19.17 ± 2.01 | 21.52 ± 3.40 |
| **Linearized Stiffness (MPa)** |  |  |  |  |
| Circumferential (Cϑϑϑϑ) | 0.08 ± 0.01 | 0.09 ± 0.01 | 0.06 ± 0.00 | 0.07 ± 0.00 |
| Axial (Czzzz) | 0.16 ± 0.03 | 0.15 ± 0.03 | 0.15 ± 0.02 | 0.17 ± 0.02 |
| **Distensibility (mmHg^-1^)** | 0.05 ± 0.00 | 0.04 ± 0.01 | 0.05 ± 0.00 | 0.05 ± 0.00 |
| **PWV (m/s)** | 1.60 ± 0.06 | 1.80 ± 0.12 | 1.63 ± 0.08 | 1.56 ± 0.02 |
| **Stored Energy (kPa)** | 5.33 ± 0.38 | 5.44 ± 0.95 | 3.71 ± 0.39 | 4.55 ± 0.27 |

**Supplemental Table S3: Proximal pulmonary artery geometry and mechanics evaluated at *in-vivo* pressures per group**

|  | **Female Control**  **n = 5** | **Male Control**  **n = 6** | **Female Silica**  **n = 5** | **Male Silica**  **n = 5** |
| --- | --- | --- | --- | --- |
| **Unloaded Dimensions** |  |  |  |  |
| Wall Thickness (μm) | 57.76 ± 3.79 | 57.54 ± 3.79 | 60.60 ± 1.30 | 61.98 ± 1.25 |
| Outer Diameter (μm) | 674.60 ± 46.57 | 675.18 ± 54.70 | 620.60 ± 35.31 | 642.20 ± 21.21 |
| Axial Length (mm) | 2.12 ± 0.12 | 1.95 ± 0.31 | 2.28 ± 0.10 | 2.33 ± 0.16 |
| **Loaded Dimensions** | **P = 15.0 mmHg** | **P = 15.0 mmHg** | **P = 25.0 mmHg** | **P = 25.0 mmHg** |
| Wall Thickness (μm) | 35.44 ± 2.39 | 34.82 ± 2.23 | 29.03 ± 1.53 | 27.98 ± 0.49 |
| Outer Diameter (μm) | 908.67 ± 54.57 | 900.90 ± 42.43 | 1090.34 ± 17.94 | 1198.78 ± 25.94 |
| Axial Length (mm) | 418.90 ± 26.79 | 415.63 ± 21.49 | 516.14 ± 9.24 | 571.41 ± 12.64 |
| *in vivo* Axial Stretch (λziv) | 1.15 ± 0.03 | 1.18 ± 0.03 | 1.10 ± 0.02 | 1.10 ± 0.02 |
| *in vivo* Circumferential Stretch (λϑiv) | 1.43 ± 0.05 | 1.43 ± 0.09 | 1.93 ± 0.12 | 2.02 ± 0.05 |
| **Cauchy Stresses (kPa)** |  |  |  |  |
| Circumferential (σϑ) | 23.95 ± 1.84 | 24.50 ± 2.33 | 60.08 ± 3.97 | 68.07 ± 1.12 |
| Axial (σz) | 23.92 ± 2.92 | 24.29 ± 3.34 | 40.27 ± 3.23 | 45.30 ± 3.52 |
| **Linearized Stiffness (MPa)** |  |  |  |  |
| Circumferential (Cϑϑϑϑ) | 0.08 ± 0.01 | 0.09 ± 0.01 | 0.25 ± 0.01 | 0.29 ± 0.01 |
| Axial (Czzzz) | 0.16 ± 0.03 | 0.15 ± 0.03 | 0.29 ± 0.07 | 0.32 ± 0.02 |
| **Distensibility (mmHg^-1^)** | 0.05 ± 0.00 | 0.04 ± 0.01 | 0.03 ± 0.00 | 0.03 ± 0.00 |
| **PWV (m/s)** | 1.60 ± 0.06 | 1.80 ± 0.12 | 2.25 ± 0.07 | 2.22 ± 0.01 |
| **Stored Energy (kPa)** | 5.33 ± 0.38 | 5.44 ± 0.95 | 14.73 ± 1.51 | 17.06 ± 0.45 |
